# Functional Complexity of Engineered Neural Networks Self-Organized on Novel 3D Interfaces

**DOI:** 10.1101/2024.06.01.596939

**Authors:** Nicolai Winter-Hjelm, Kasper Grøndahl Klausen, Amund Stensrud Normann, Axel Sandvig, Ioanna Sandvig, Pawel Sikorski

**Author notes:** Correspondence, &. These authors have contributed equally to this work (Ioanna Sandvig, Pawel Sikorski).

## Abstract

Engineered neural networks are indispensable tools for studying neural function and dysfunction in controlled microenvironments. *In vitro*, neurons self-organize into complex assemblies with structural and functional similarities to *in vivo* circuits. Traditionally, these models are established on planar interfaces, but studies suggest that the lack of a three-dimensional growth space affects neuronal organization and function. While methods supporting 3D growth exist, reproducible 3D neuroengineering techniques compatible with electrophysiological recording methods are still needed. In this study, we developed a novel biocompatible interface made of the polymer SU-8 to support 3D network development. Using electron microscopy and immunocytochemistry, we show that neurons utilize these 3D scaffolds to self-assemble into complex, multi-layered networks. Furthermore, interfacing the scaffolds with custom microelectrode arrays enabled characterizing of electrophysiological activity. Both planar control networks and 3D networks displayed complex interactions with integrated and segregated functional dynamics. However, control networks showed stronger functional interconnections, higher entropy, and increased firing rates. In summary, our interfaces provide a versatile approach for supporting neural networks with a 3D growth environment, compatible with assorted electrophysiology and imaging techniques. This system can offer new insights into the impact of 3D topologies on neural network organization and function.

## Introduction

In the developing brain, neurons self-organize over time into networks with high computational capacity. This organization takes place under the influence of the dynamic interplay between intrinsic neuronal self-organizing properties, spatiotemporally regulated chemical and structural guidance cues, and experience-dependent plasticity (1, 2). Engineered neural networks can be designed to recapitulate key hallmarks of this structural and functional complexity *in vitro* (3– 5). Such model systems facilitate studies of neural network function and dysfunction at the micro- and mesoscale level in a highly controllable microenvironment (6, 7). Typically, these networks are established on planar interfaces. Studies however indicate that physical dimensionality, i.e., planar versus 3D configuration, can influence cellular characteristics such as morphology, proliferation rates, differentiation, migration, gene expression, electrical activity, and susceptibility to pathology (8–12). However, whether and to what extent physical dimensionality may impact the networks’ functional dynamics remains largely under-explored. Addressing these questions requires accessible and reproducible methods for modulating the physical dimensionality of the networks. Furthermore, such methodologies need to be compatible with electrophysiology and imaging tools for advanced characterization of the networks’ structure-function dynamics. Advancements in the relevant technologies can yield new insights into the relationship between neural network topology and function, and also expand the repertoire of techniques available for preclinical disease modelling and drug screening (13–16).

In recent years, a range of methodologies have been developed to provide neurons with a 3D microenvironment in which they can grow and self-organize (17). One widely applied technique is the use of hydrogels, which mimic key properties of the extracellular matrix found in tissue (18). However, hydrogels can complicate cell handling, nutrient and waste transport, cell culture maintenance, and electrophysiological characterization. The latter issue arises largely due to the lack of robust 3D electrode technologies, which leads to severe subsampling in the networks as only the activity of cells in the bottom layer can be recorded (19). As an alternative approach, allowing the cells to self-organize in 3D scaffolds may be a good alternative to full scale 3D cultures as the accessibility to the entirety of the neural network remains (17). Scaffold-based 3D interfaces have been shown to support more metabolically and computationally efficient network organization without complicating experimental procedures compared to traditional planar models (20, 21). Over the past decade, various scaffold structures have been utilized, including pillars (22–24), cylinders (25), steps and grooves (26), and polymer structures with heterogeneous morphologies (27, 28). While these approaches represent incremental progress, they often remain too complex to reproduce and utilize effectively.

Neural network topology and function are closely linked, with topology shaping network function and network activity influencing topology (29–31). Although several studies have shown that 3D topologies support altered morphological characteristics, only a limited number of studies have conducted functional characterization of engineered networks grown in 3D scaffolds (28, 32, 33). These studies found that 3D scaffolds promoted a more dynamical regime between high synchrony and more localized activity in the neural networks than planar interfaces (33). Additionally, while 3D networks exhibited periods of highly synchronized bursts, the frequency and duration differed from those on 2D interfaces (32). Moreover, 3D networks showed intermittent periods of desynchronized spiking activity, resembling what can be seen *in vivo*. Electrical stimulations of the 3D networks were also reported to induce more discontinuous spreading of evoked activity compared to networks grown on planar interfaces (32). These findings highlight the applicability and significance of utilizing geometrical scaffolds to manipulate and study the impact of physical dimensionality on the functional dynamics of neural networks.

Inclined lithography, first introduced by Han *et al*. in 2004 (34), is a relatively new approach for creating 3D structures. In 2018, Li *et al*. demonstrated its utility for constructing 3D lattices for neural growth (35). They applied this technique to create SU-8 molds, which were later replica-molded into PDMS structures. Their method showed great promise for structuring 3D networks with high reproducibility and ease. Yet, a limitation with this approach is the challenge of interfacing the PDMS structures with microelectrode arrays (MEAs) to facilitate electrophysiological recordings with high temporal resolution. In this study, we show that an approach in which the 3D scaffolds are made directly in SU-8 supports integration with MEAs. SU-8 is an epoxy-based photoresist widely applied in microfabrication of biomedical devices (36, 37). To our knowledge, only one comparable study has explored the use of SU-8 scaffolds for structuring 3D neural networks in the past (38). Nevertheless, the fabrication approach used in this study would be challenging to adapt for most labs, and require further optimization for integration with MEAs. In this study, we show that combining inclined SU-8 scaffolds with nanoporous microelectrode arrays (5) ensures high consistency and reproducibility, proper coupling between the networks and the electrodes, and supports neural outgrowth and 3D network formation. We furthermore show that these 3D scaffolds facilitate the establishment of neural networks with high computational capacity and distinct functional characteristics compared to networks established on planar interfaces.

## Materials and Methods

### Design of Experimental Interface

Designs for microfab-rication were created in Clewin 4 (WieWeb Software, Enschede) and are shown in **Figure S1**. The MEAs were designed to be compatible with the MEA2100 workstation from Multichannel Systems, featuring 59 electrodes with diameters of either 50 µm (control-MEAs) or 100 µm (tripod-MEAs) evenly distributed across the chip. Additionally, a 1.5 mm diameter reference electrode was positioned at the chip’s periphery. To create the tripod structures, photomasks with 15 µm apertures arranged in a hexagonal pattern with 50 µm interspacing were fabricated. The hexagonal pattern was used to create overlapping tripod structures. Three apertures were removed at the position of each electrode to avoid tripods covering up the electrodes. 50 µm diameter clamping lines were included to anchor the scaffolds to the substrate and prevent delamination. 4 clamping lines ran through the central area of the design, while another clamping line was positioned along the circumference. The diameter of the SU-8 pattern was set to 5.0 mm to fit within 6.0 mm diameter PDMS cell culturing chambers. 6 control-MEAs and 5 tripod-MEAs were used for electrophysiological experiments.

### Fabrication of Microdevices

#### Microelectrode Arrays

The protocol for fabricating micro-electrode arrays was adapted from our previous work (5, 6), with modifications to integrate SU-8 3D structures. 1 mm thick, 4-inch borosilicate wafers (100 mm Borofloat33, Plan Optik) were used as substrates. These wafers were cleaned in acetone and subsequently IPA for 1 min each to remove organic contaminants, then plasma cleaned for 5 min in 100 sccm O_2_ plasma at 20 kHz generator frequency (Femto Plasma Cleaner, Diener Electronics). A 2 min dehydration bake was conducted at 100 *◦*C to remove residual moisture.

The photoresist ma-N 440 (micro resist Technology GmbH) was spin coated onto the substrates at 3000 rpm for 42 s at 500 rpm/min, achieving a final thickness of 4 µm (spin150, SPS-Europe B.V.). After a 10 min rest, the film was soft baked at 95 *◦*C for 5 min. The MEA design was transferred onto the resist using a maskless aligner (MLA150, Heidelberg) with a 405 nm laser at 1800 mJ/cm_2_. After exposure, the resist was developed in ma-D332/s (micro resist Technology GmbH) for 90±10 s, then rinsed thoroughly in DI water. Before e-beam evaporation, the substrates were descummed in 100 sccm O_2_ plasma for 1 min at 20 kHz. An adhesive layer of 50 nm titanium was evaporated onto the substrates at 5 Ås_*−*1_, followed by 100 nm platinum at 2 Ås_*−*1_ (E-beam Evaporator, Pfeiffer Vacuum Classic 500). Finally, lift-off was performed using acetone, followed by an IPA rinse. Before depositing the passivation layer, another 1 min descum step was conducted. Subsequently, a 470 nm thick insulation layer of silicon nitride (Si_3_N_4_) was deposited onto the substrates via plasma-enhanced chemical vapour deposition at 300 *◦*C for 30 min. This process utilized a gas mixture consisting of 20.0 sccm SiH_4_, 20.0 sccm NH_3_ and 980 sccm N_2_ (PlasmaLab System 100-PECVD, Oxford Instruments).

Next, a second lithographic step using ma-N 440, following the same protocol as previously described, was used to define an etch mask. Inductively coupled plasma with a gas composition of 50.0 sccm CHF_3_, 10.0 sccm CF_4_ and 7.0 sccm O_2_ was subsequently employed for 6.5 min to dry etch the silicon nitride above the electrodes and contact pads (Plasmalab System 100 ICP-RIE 180, Oxford Instruments).

After etching, the substrates to be interfaced with tripod structures were diced into 49×49 mm square MEAs using a wafer saw (DAD323, DISCO). Substrates designated for control-MEAs were diced after conducting electrodeposition of platinum black (further described below). The ma-N 440 was then removed in acetone, followed by a rinse in IPA. To remove hardened photoresist and oxidize the top nanometres of the silicon nitride layer into silicon dioxide (SiO_2_), the surface underwent plasma cleaning for 10 min in 160 sccm O_2_ at 32 kHz generator frequency.

#### Mask Fabrication for Inclined Lithography

For detailed instructions on the mask fabrication, please refer to the supplementary materials. Illustrative steps of the mask fabrication process are depicted in **Figure S2. *SU-8 Tripod Structures***. An illustration of the fabrication process of SU-8 tripod structures can be seen in **Figure 1A**. Tripods were either fabricated directly on #1.5 glass coverslips (24×24 mm Menzel-Gläser, VWR International) precleaned with acetone, IPA and oxygen plasma, or aligned onto MEAs. First, a dehydration bake was conducted at 150 *◦*C for 6 min. Subsequently, to enhance SU-8 adhesion, the substrates were primed with 1,1,1,3,3,3-Hexamethyldisilazane (Acros Organics, 430851000) in a dessicator for 3 h.

**Figure 1.**
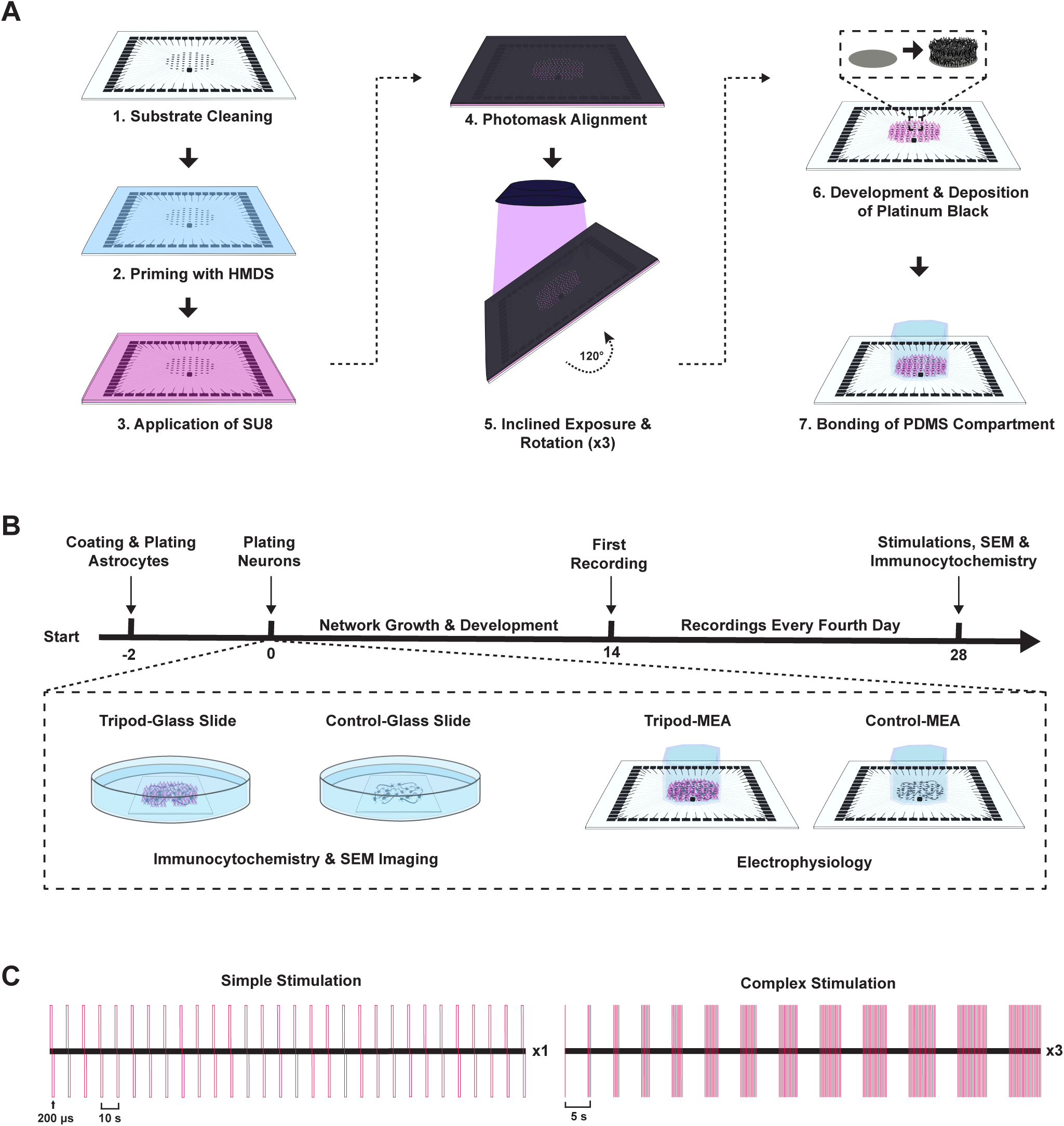
Experimental setup. **(A.)** Illustration of the fabrication process for creating tripod structures on a microelectrode array (MEA) using inclined photolithography.**(B.)** Timeline for cell experiments. Astrocytes and neurons were cultured on MEAs or glass coverslips with or without tripod structures. Electrophysiological recordings were conducted every fourth day from day 10 to day 28 *in vitro* (DIV). **(C.)** Network stimulations were conducted following the recording of spontaneous activity at 28 DIV. First, a simple stimulation protocol consisting of a train of 30 consecutive spikes with a 10 s interspike interval was delivered to the most active electrode on the MEA. Secondly, an advanced stimulation protocol with a linearly increasing number of pulses (1:2:25), each with 5 s intervals between trains, was delivered to three highly active electrodes evenly spaced at the periphery of the MEA. The complex stimulation protocol was repeated three times. The illustrations are not drawn to scale.

Before applying the photoresist, the substrate underwent another dehydration baked at 150 *◦*C for 6 min. Subsequently, the photoresist SU-8 3050 (Kayaku Advanced Materials, Y311075 0500L1GL) was spin coated onto the substrate in two steps: First at 3000 rpm for 40 s with a ramp-up speed of 300 rpm/min, and then at 4000 rpm for 5 s with a ramp-up speed of 1000 rpm/min, resulting in a final thickness of 45 µm (spin150, SPS-Europe B.V.). After spin coating, the substrate was allowed to rehydrate at room temperature (RT) for 1 h prior to soft bake. The soft bake was performed using a programmable hot plate (HP8, Süss MicroTec), starting at 30 *◦*C for 5 min, followed by a gradual increase in temperature to 95 *◦*C at a rate of 2 *◦*Cmin_*−*1_, where it was kept for 35 min. Subsequently, the substrate cooled down on the hotplate at a rate of 1 *◦*Cmin_*−*1_.

The photomask was aligned onto the MEA using custom-made alignment marks (see **Figure S1B**) under a stereomicroscope (Nikon SMZ800), the the chromium layer facing down, and secured to the sample using scotch tape. Subsequently, the substrate was mounted at the center of a custom-made exposure stage. The stage was designed in Tinker-CAD and 3D printed using an Ultimaker 2+ (Ultimaker) with a polyactic acid filament (3DNet, PLA 2.85), and using a 0.6 mm nozzle at 230 *◦*C and a build plate temperature of 60 *◦*C. The design can be seen in **Figure S3A**, and an image of the full exposure setup in **Figure S3B**. The stage had a 45*◦* tilt when mounted to the tower, featuring an equilateral triangle at the back to facilitate three consecutive exposures with 120*◦* rotations. For UV curing, a 10 W UV lamp equipped with eight 365 nm uncollimated UV LED diodes arranged in a 2×4 array (XT004, Extral Light Store) was utilized. The radiation intensity was quantified at 21 mW/cm_2_ at a distance of 5 cm using a UV-Optometer from SÜSS MicroTec. To shield photoresist not covered by the photomask from exposure, aluminium foil served as a shadow mask, as demonstrated in **Figure S3C**. Finally, an opaque enclosure was employed to shield the setup from ambient light during exposures. Each sample underwent three exposures, with a 120*◦* rotation of the stage between exposures, as visualized in **Figure S3D**. The UV-lamp was activated for 25 s per exposure, resulting in a dosage of 525 mW/cm_2_.

Following exposure, the substrate underwent a post-exposure bake, commencing at 30 *◦*C for 5 min before gradually ramping up the temperature to 60 *◦*C at 1 *◦*Cmin_*−*1_, where it was kept for 60 min. Subsequently, the substrate was allowed to cool gradually on the hotplate at a rate of 1 *◦*Cmin_*−*1_. Following this, development was performed using mr-Dev 600 (micro resist Technology GmbH) for 7 min, stirring the developer every 30 s and dousing the substrate with fresh developer every 2 min. Afterward, a thorough rinse in IPA wad administered, and the sample carefully dried using a nitrogen gun. A hard bake followed, starting at 30 *◦*C for 2 min before gradually increasing the temperature to 180 *◦*C at a rate of 7.5 *◦*Cmin_*−*1_, where it was kept for 60 min. The substrate was then allowed to cool on the hotplate at a rate of 2 *◦*Cmin_*−*1_.

#### Deposition of Nanoporous Platinum

A thin layer of platinum black was electrodeposited onto the electrodes using a PP-Type Wafer Plating Electroplating Laboratory System from Yamamoto. For the control-MEAs, electrodeposition was conducted prior to dicing the substrates into 49×49 mm substrates. This method enabled the larger metal pads at the periphery of the design to connect up the smaller contact pads during depositions, allowing for concurrent electrodeposition of up to 15 electrodes. These connections were subsequently severed upon dicing the wafer, as depicted in **Figure S1A**. For the tripod-MEAs, electrodeposition took place after dicing the substrates. To facilitate the simultaneous deposition of multiple electrodes, conductive copper tape was utilized to interconnect the contact pads during depositions. The copper tape was then linked to the potentiostat using an alligator clip.

The electrolyte bath was comprised of an aqueous solution containing 2.5 mmol chloroplatinic acid (H_2_PtCl_6_, 8 wt% H_2_O, Sigma-Aldrich, 262587). The wafers were partially submerged in this solution using a custom-built wafer holder, ensuring that liquid covered only the electrodes and not the contact pads, as described in our previous work (5). A Red Rod REF201 Ag/AgCl electrode (Hatch) served as the reference electrode, while a platinized titanium plate functioned as the counter electrode. To minimize diffusion thickness, a paddle agitator was operated at 60.0 rpm, while the temperature was maintained at a constant 30 *◦*C using an external temperature controller. Electrodeposition was conducted using a Palmsens 4 potentiostat (Palmsens) in chronoamperometric mode, applying a constant voltage of −0.4 V for a duration of 3 min for all depositions.

#### Biocompatibility Treatment

To ensure the tripod structures were hydrophilic and biocompatible, the substrates underwent plasma treatment in 200 sccm O_2_ plasma for 1 min at 40 kHz generator frequency. Subsequently, they were rinsed in ultrasonically agitated IPA for 15 min to draw out potentially toxic leachants.

#### PDMS Compartments

A 4-inch silicon wafer (Siltronix) was washed consecutively in acetone and isopropanol (IPA) for 1 min each to eliminate organic contaminants. Following this, a 5 min plasma clean was conducted using 100 sccm O_2_ plasma at a generator frequency of 20 kHz (Femto Plasma Cleaner, Diener Electronics). Subsequently, a 5 min dehydration bake was performed at 120 *◦*C to remove moisture. Silicon elastomer and curing agent (SYLGARD®184 elastomer kit, Dow Corning) were mixed, degassed, and cast atop the silicon wafer at a 10:1 ratio. The PDMS was cured in an oven (TS8056, Termaks) at 65 *◦*C for 4 h. Post-curing, the PDMS was peeled from the mould, and cell compartments were cut out with a 6 mm diameter puncher. PDMS residues were removed using scotch tape, and the chips subsequently washed in acetone, 96 % ethanol and DI water for 1 min each. Finally, the chips were left to dry overnight.

#### Assembly of Tripod-MEA Interfaces

The PDMS chips were irreversibly bonded to nanoporous MEAs, with or without tripod structures. Both the PDMS device and MEA underwent plasma treatment with O_2_ plasma for 1 min at a flow rate of 200 sccm O_2_ and a generator frequency of 40 kHz. Directly after, the PDMS chip was gently pressed onto the MEA. To aid alignment of the PDMS chip to the MEA, two drops of 70 % ethanol were placed between the PDMS chip and the MEA, with alignment performed under a stereomicroscope.

Bonding was finalized on a hotplate at 100 *◦*C for 1 min followed by a 5 min rest at RT under gentle pressure. To remove residual ethanol, three consecutive washes in DI water were conducted at 10 min intervals. Finally, the chips were filled with DI water to maintain hydrophilicity and sterilized overnight under UV light in a biosafety cabinet.

### Cell Culturing and Staining

#### Coating of Culturing Interfaces

A timeline for the cell experiments is depicted in **Figure 1B**. Following sterilization and preceding surface coating, the samples were immersed in DMEM, low glucose (Gibco™, 11885084) for a minimum of 48 h to remove any potential toxic leachants remaining in the SU-8 structures. Subsequently, the cell medium was replaced with a 0.1 mg/mL Poly-L-Ornithine solution (PLO) (Sigma-Aldrich, A-004-C), and the chips were incubated overnight in a fridge at 4 *◦*C. The following day, the chambers underwent three washes with MQ water before being filled with a laminin solution containing 16 µg/mL natural mouse laminin (Gibco™, 23017015) in phosphate-buffered saline (PBS, Sigma-Aldrich, D8537). Subsequently, the chips were incubated at 37 *◦*C, 5 % CO_2_ for 2 h.

#### Cell Seeding and Maintenance

After coating, the laminin solution was substituted with pre-warmed astrocyte medium comprising DMEM low glucose supplemented with 15 % Fetal Bovine Serum (Sigma-Aldrich, F9665), and 2 % Penicillin-Streptomycin (Sigma-Aldrich, P4333). Rat astrocytes (Gibco™, N7745100) were then plated at a density of 100 cells/mm_2_, equivalent to 4000 cells per culturing chamber. Following two days of expansion, the astrocyte medium was replaced with neuronal medium composed of Neurobasal Plus Medium (Gibco™, A3582801) supplemented with 2 % B27 Plus (Gibco™, A358201), 1 % Gluta-Max (Gibco™, 35050038), and 2 % Penicillin-Streptomycin (Sigma-Aldrich, P4333). Additionally, Rock Inhibitor (Y-27632 dihydrochloride, Y0503, Sigma-Aldrich) was included at a concentration of 0.1 % during plating to enhance survival rates. Rat cortical neurons from Sprague Dawley rats (Gibco, A36511) were plated at a density of 1000 cells/mm_2_, equalling 40,000 cells per chamber. Half of the cell medium was replaced with fresh medium 4 h after plating, and again after 24 h. Subsequently, half of the cell medium was replaced every other day until the cultures were terminated at 28 DIV. To mitigate biases arising from batch-to-batch variability, all astrocytes and neurons originated from the same batch and cell vials.

#### Immunocytochemistry

For immunocytochemistry (ICC), cells were plated onto glass coverslips with or without tripod structures, positioned in multiwell plates. The cells were fixed using glyoxal solution based on the protocol by Richter *et al*.(39). The fixative comprised 71 % MQ water, 20 % absolute ethanol (Kemetyl, 100 %), 8.0 % Glyoxal solution (Sigma-Aldrich, 128465), and 1 % acetic acid (Sigma-Aldrich, 1.00063). Following incubation in the fixative at RT for 15 min, the chambers were washed three times with phosphate-buffered saline (PBS, Sigma-Aldrich, D8537) for 5 min each. Next, the cells were permeabilized with 0.5 % Triton-X (Sigma-Aldrich, 1086431000) diluted in PBS for 5 min, followed by two additional washes in PBS for 5 min each. Subsequently, the cells were incubated in blocking solution containing 5 % goat serum (Abcam, ab7481) diluted in PBS on a shaker table (30 rpm) at RT for 1 h. Thereafter, primary antibodies mixed with 5 % goat serum in PBS were added to the chambers, and the cells incubated on a shaker table at 30 rpm at 4 *◦*C overnight. An overview of the primary antibodies can be found in **Table 1**.

**Table 1.**
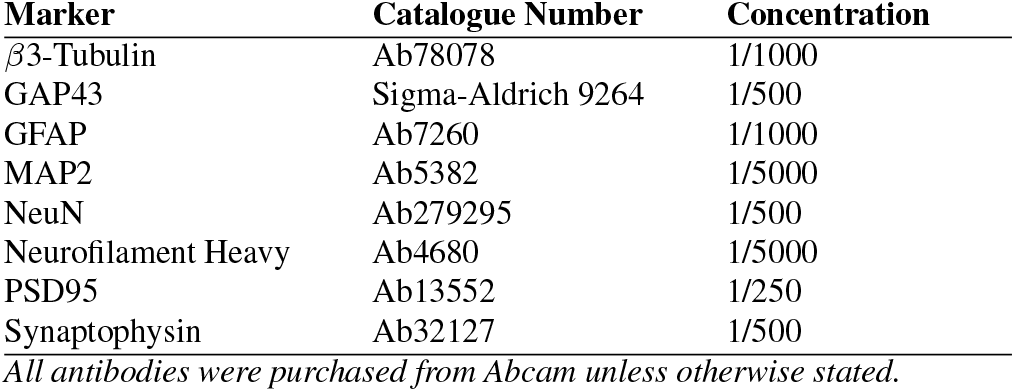
Antibodies and concentrations used for Immunocytochemistry.

| Marker | Catalogue Number | Concentration |
| --- | --- | --- |
| β3-Tubulin | Ab78078 | 1/1000 |
| GAP43 | Sigma-Aldrich 9264 | 1/500 |
| GFAP | Ab7260 | 1/1000 |
| MAP2 | Ab5382 | 1/5000 |
| NeuN | Ab279295 | 1/500 |
| Neurofilament Heavy | Ab4680 | 1/5000 |
| PSD95 | Ab13552 | 1/250 |
| Synaptophysin | Ab32127 | 1/500 |

All antibodies were purchased from Abcam unless otherwise stated.

The following day, the chambers were washed three times with PBS for 5 min each. Subsequently, the cells were incubated in a secondary antibody solution consisting of 0.2 % secondaries and 5 % goat serum diluted in PBS on a shaker table (30 rpm) at RT in the dark for 3 h. Hoechst nuclear stain (Abcam, ab228550) was then added at a concentration of 0.2 % (diluted in PBS), and the cells were incubated for an additional 30 min on the shaker table. Following this, the chips were washed three more times in PBS, followed by two final washes in MQ water for 5 min each.

Imaging was performed using a Zeiss 800 Airyscan Confocal Laser Scanning Microscope (CLSM) with Zeiss Zen Blue software. The microscope was equipped with a LSM 800 Laser Module URGB, featuring diode lasers at wavelengths of 405 nm (5 mW), 488 nm (10 mW), 561 nm (10 mW) and 640 nm (5 mW). A Zeiss Axiocam 503 color camera, 2.8 Mpix, 38 fps (color) was furthermore used for image acquisition.

#### Sample Dehydration and SEM Imaging

The samples underwent two rinses in pre-warmed PBS to remove dead cells and debris. Fixation was achieved using 2.5 % glutaraldehyde (G5882, Sigma-Aldrich) in Sorensen’s phosphate buffer (0.1 M, pH 7.2). The fixative was thawed overnight in a refrigerator and left in a fume hood for 1 h prior to fixation to equilibrate to RT. Any precipitates were carefully resuspended through agitation with a micropipette. The cells were then immersed in the fixative for 2 h at RT before being transferred to a refrigerator at 4 *◦*C overnight. Following fixation, water was chemically extracted from the specimens via serial dehydration in 25 %, 50 %, 70 %, 80 %, 90 % and 100 % ethanol, with each step lasting 5 min. The final ethanol dehydration step was repeated twice to ensure complete removal of water. Subsequently, a Leica EM CPD300 Critical Point Dryer was employed to replace the ethanol with CO_2_, with the instrument set to a 120 s delay time, exchange speed of 1, with 18 cycles and gas out speed of 50 % (slow). After drying, a Cressington 208 HR B Sputter Coater was used to deposit 15 nm of platinum/palladium onto the samples. During coating, the samples were tilted in a cyclic manner from −45*◦* to 45*◦* with a period of 20 s. Imaging was performed using an APREO Field Emission Scanning Electron Microscope from FEI with an EDT detector to capture secondary electrons. The beam current ranged between 25 and 50 pA, and the acceleration voltage was set between 4.0 and 10.0 kV.

### Electrophysiology and Data Analysis

#### Electrophysiological Recordings

Recordings were conducted using a MEA2100 workstation (Multichannel Systems) with a sampling rate of 25000 Hz. A temperature controller (TC01, Multichannel Systems) maintained the temperature at 37 *◦*C. To ensure sterility during recordings, a 3D-printed plastic construct covered by a gas-permeable membrane was used. Prior to each recording, the networks were allowed to equilibrate for 5 min at the recording stage. Each recording session lasted 15 min and was always conducted the day following media changes.

#### Electrical Stimulations

Stimulations were performed following the recordings of spontaneous network activity on 28 DIV using two different patterns, referred to as “simple” or “complex”. The simple stimulation protocol consisted of a train of 30 consecutive stimulation pulses at ±800 mV amplitude (positive phase first) with a 200 µs duration and a 10 s interspike interval. This spike train was delivered to the electrode with the highest firing rate (spikes/second), indicating pacemaker activity, in accordance with previous literature (40–43). The complex stimulation protocol consisted of sequences with a linearly increasing number of stimulation pulses (1:2:25), each with ±800 mV amplitude (positive phase first), 200 µs pulse duration, and 200 µs interspike interval, and each sequence was delivered with a 5 s interval. This pattern was repeated three times and delivered to three highly active electrodes distributed at the periphery of the MEA. The two stimulation protocols are illustrated in **Figure 1C**.

#### Data Analysis

All data analysis, including filtering and spike detection, was conducted in Matlab R2021b. Custom colormaps were created using the Matlab functions *linspecer* (44) and *colorBrewer2* (45), based on the web tool colorBrewer developed by Brewer *et al*. (46).

Raw data was filtered using a 4th-order Butterworth bandpass filter to remove high-frequency noise (above 3000 Hz) and low-frequency fluctuations (below 300 Hz). A notch filter eliminated 50 Hz noise from the power supply mains. Both filters employed zero-phase digital filtering via the Matlab function *filtfilt* to preserve the relative position of the detected spikes. Spike detection was conducted using the Precise Timing Spike Detection (PTSD) algorithm developed by Maccione *et al*. (47). The threshold was set to 9 times the standard deviation of the noise, with a maximum peak duration of 1 ms and a refractory time of 1.6 ms. Raster plots were generated using a customized version of the *SpikeRasterPlot* function developed by Kraus (48).

Burst detection was performed using the logISI method developed by Pasquale *et al*. (49). A minimum of 4 consecutive spikes was required for burst detection, with a hard threshold for the interspike interval set to 100 ms both within and outside of the burst core. This threshold was chosen based on the original manuscript reporting the method (49), as well as manual inspection of the burst detection performance. For analyzing spontaneous network activity, network bursts were identified using the logIBEI approach (50). At least 20 % of active electrodes in the network needed to participate for the activity to be classified as a network burst. Network synchrony was measured using the coherence index (51).

The data was binned into 20 ms intervals. To estimate the casual relationship between activity recorded by the electrodes, mutual information was calculated between all electrode pairs using the information theory toolbox by Timme & Lapish (52). Entropy and complexity calculations were conducted with the toolbox developed by Marshall *et al*. (53), based on the complexity measure introduced by Tononi *et al*. (29). For detailed methodologies, refer to the original papers (29, 53), but key metrics used in this study are briefly introduced here. Network entropy was calculated using **Equation 1**:

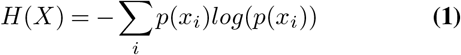

where *x*_*i*_ is defined as the joint state of all electrodes in a given time bin, and the probability of that state *p*(*x*_*i*_) is the number of occurrences of that state divided by the total number of time bins. Total correlation, also referred to as network integration, was calculated as the difference between the joint entropy of a set *j* of *k* neurons and their individual entropies, using **Equation 2**:

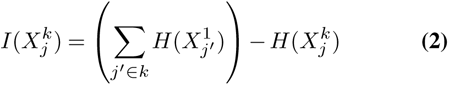

Network complexity is designed to favor a balance between functional segregation (i.e. statistical independence) and functional integration (i.e. deviations from such statistical independence). This measure was calculated using **Equation 3**:

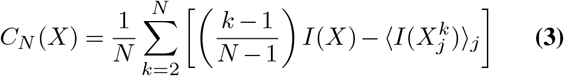

Given the high number of possible permutations in the system, 100 permutations were randomly selected for calculating the average integration in the complexity calculations.

For graph theoretical analysis, the functional connectivity matrices were thresholded to include only the most significant, direct connections. First, the binned matrices were shuffled, and mutual information was recalculated to estimate the maximum functional connectivity between two electrodes by chance. Connections weaker than this threshold were removed. Additionally, only the 10 % strongest connections were retained for the graph analysis. Various thresholds ranging from 5 % to 100 % were tested, which did not significantly impact the graph analysis outcome.

Graph theoretical metrics were calculated using the Brain Connectivity Toolbox developed by Rubinov & Sporns (54). These metrics were used to create custom-designed graphs where node colour represents spike frequency (spikes/s), node size indicates PageRank centrality, edge colour shows functional connectivity (pairwise mutual information), and node edge colour indicates community belonging calculated using the Louvain algorithm (55). Nodes were positioned according to the relative position of the electrodes on the MEAs. Small world propensity was calculated using the method developed by Muldoon *et al*. (56).

Stimulation data was run through the SALPA filter developed by Wagenaar *et al*. (57). Each stimulation time point was followed by a 15 ms blanking period to prevent stimulation artifacts from being detected as spikes. Peristimulus time histograms (PSTHs) were generated for each MEA by binning the data in the 300 ms following a stimulation into 20 ms intervals. The average response of the network in each bin for the 60 individual stimulations was plotted.

#### Statistical Analysis

For longitudinal data analysis, Generalized Linear Mixed-Effect Models (GLMMs) were constructed using SPSS version 29.0.0.0. The network type (control or tripod) served as the fixed effect, while network age was considered a random effect. The network feature of interest was furthermore used as a target. Data fitting employed a gamma distribution with a log link function, chosen based on the Akaike information criterion. Multiple comparisons were adjusted using Sequential Bonferroni correction. Intergroup differences at specific time points were assessed using a two-sided Wilcoxon rank sum test in Matlab 2021b.

## Results

### Inclined Lithography is a Versatile and Reproducible Approach for Creating 3D Scaffolds of SU-8

The primary objective of this study was to establish a reproducible methodology for fabricating 3D scaffolds with inclined features on which neurons could grow and connect into three-dimensional networks. Tripods with varying diameters and densities were fabricated within 45 µm thick SU-8 layers by adjusting the size of the apertures of the photomask and their interspacing (**Figures 2A-2C**). Structural collapse was commonly observed with diameters less than 10 µm (**Figure 2A**). To achieve porous yet robust structures suitable for neuronal growth between the tripod legs, a tripod size of 15 µm diameter and 50 µm interspacing was eventually selected for all the cell experiments presented in this study (**Figures 2D** and **2E**).

**Figure 2.**
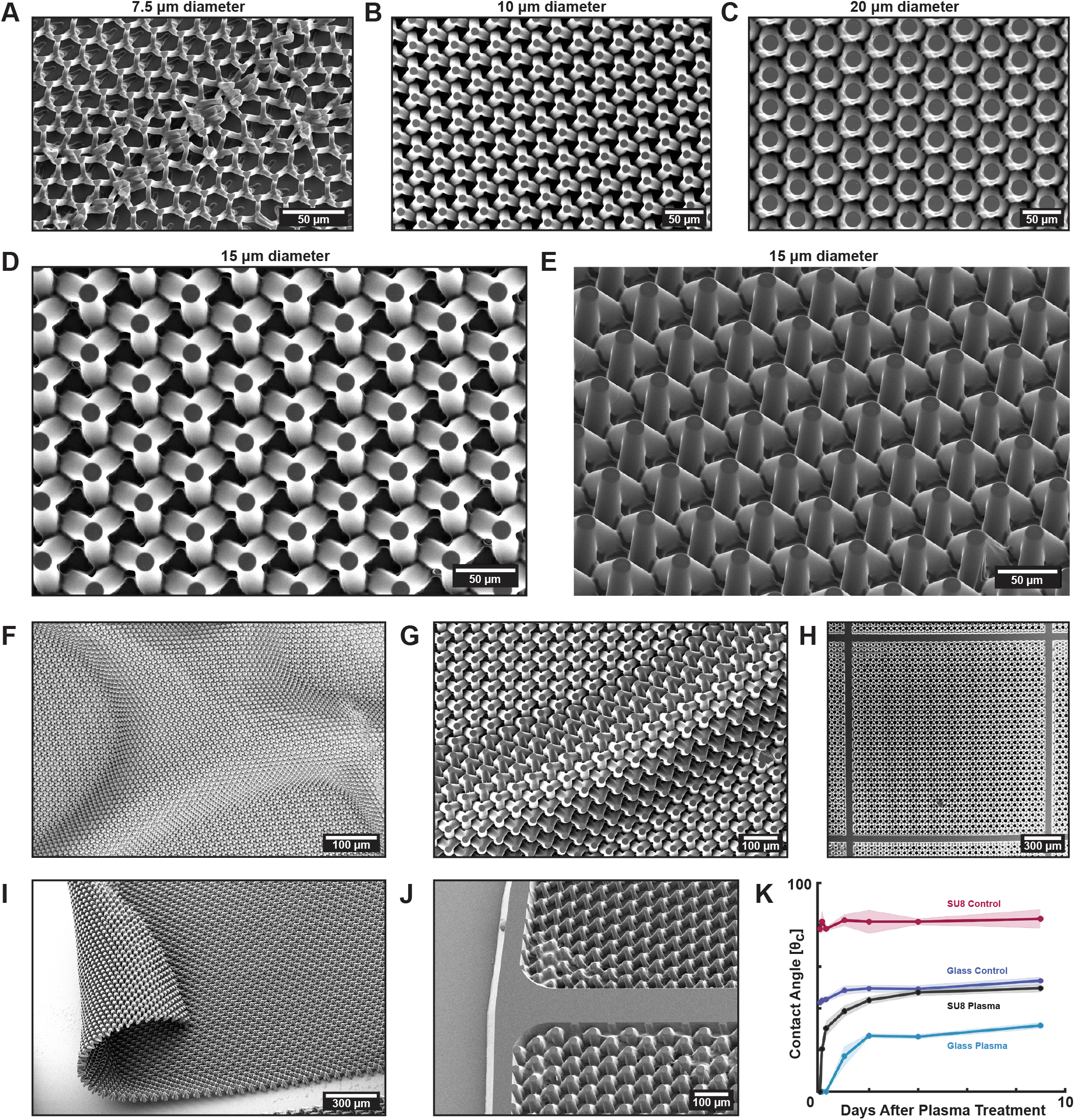
Fabrication of tripod scaffolds. **(A.)**-**(C.)** Tripods with various diameters were fabricated. Tripods with legs smaller than 10 µm commonly resulted in collapse, while those with 20 µm diameter legs or larger created densely packed arrays, limiting space for neural network growth between the structures. **(D.)**-**(E.)** Tripods with 15 µm diameter legs and 50 µm interspacing were selected for cell experiments. **(F.)**-**(G.)** Issues with heat folding and delamination of the tripod arrays arose due to the small contact area between the tripods and substrate, exacerbated by rapid temperature changes during heating. **(H.)** Introducing 50 µm diameter clamping lines of plane SU-8 at regular intervals across the arrays prevented delamination. **(I.)** Delamination also occurred along the edges of the SU-8 tripod arrays. **(J.)** Additional clamping lines along the edges ensured tripod attachment to the substrate. **(K.)** Plasma treatment of the SU-8 enhanced biocompatibility over an extended period of time by rendering it comparably hydrophilic to glass.

A reccurring challenge encountered during the process development and optimization was the delamination of tripod scaffolds from the borosilicate substrate (**Figures 2F** and **2G**). Delamination consistently became evident during the development step, likely caused by temperature fluctuations during the post exposure bake (PEB). To address this issue, planar stripes of 50 µm wide SU-8 were strategically introduced at regular intervals between the tripods, serving as clamping lines to secure the SU-8 to the substrate (**Figure 2H**). The same solution was also found effective at mitigating delamination along the edges of the tripod arrays (**Figures 2I** and **2J**). Although these clamping lines rendered parts of the available growth area for the cells planar, the proportion of the surface taken up by these lines was still considered acceptably small to not have any impact on the overall three-dimensionality of the networks.

Initial attempts to culture neurons on both planar layers of SU-8 and on tripod scaffolds resulted in significant cell death. To enhance cell viability, a series of post-processing treatments of the SU-8 were tested before cell culturing to render the surfaces more hydrophilic and remove potentially toxic leachants such as non-crosslinked resist or antimony generated from the plasma etching steps (58–60). Firstly, a hard bake of at least 60 min at 180 *◦*C was found to substantially improve cell viability, likely by facilitating proper photoresist cross-linking. Additionally, plasma cleaning of the substrates for at least 1 min rendered the hydrophilicity of the SU-8 comparably low to that of glass (**Figure 2K**). Ultimately, washing the surfaces in IPA in an ultrasonic bath for at least 15 min and subsequent incubation at 37 *◦*C in cell medium for 48 h significantly enhanced cell viability, achieving levels comparable to neurons growing on glass interfaces.

### Neurons Readily Form Multilayered Networks Within the 3D Scaffolds

Electron microscopy confirmed that neurons effectively utilized the scaffolds to self-organize into dense multilayered networks both on top of and between the tripod structures (**Figures 3B-3C**). Notably, neuronal somata were observed extending between the tripods, solely connected to the scaffolds via their neurites (**Figure 3D**). Neurites also extended between the tripods, facilitating shortcuts between neural clusters attached to the tripods (**Figure 3E**). Additionally, fasciculated axonal bundles, measuring up to a millimeter in length, were observed extending across significant portions of the arrays (**Figure 3F**).

**Figure 3.**
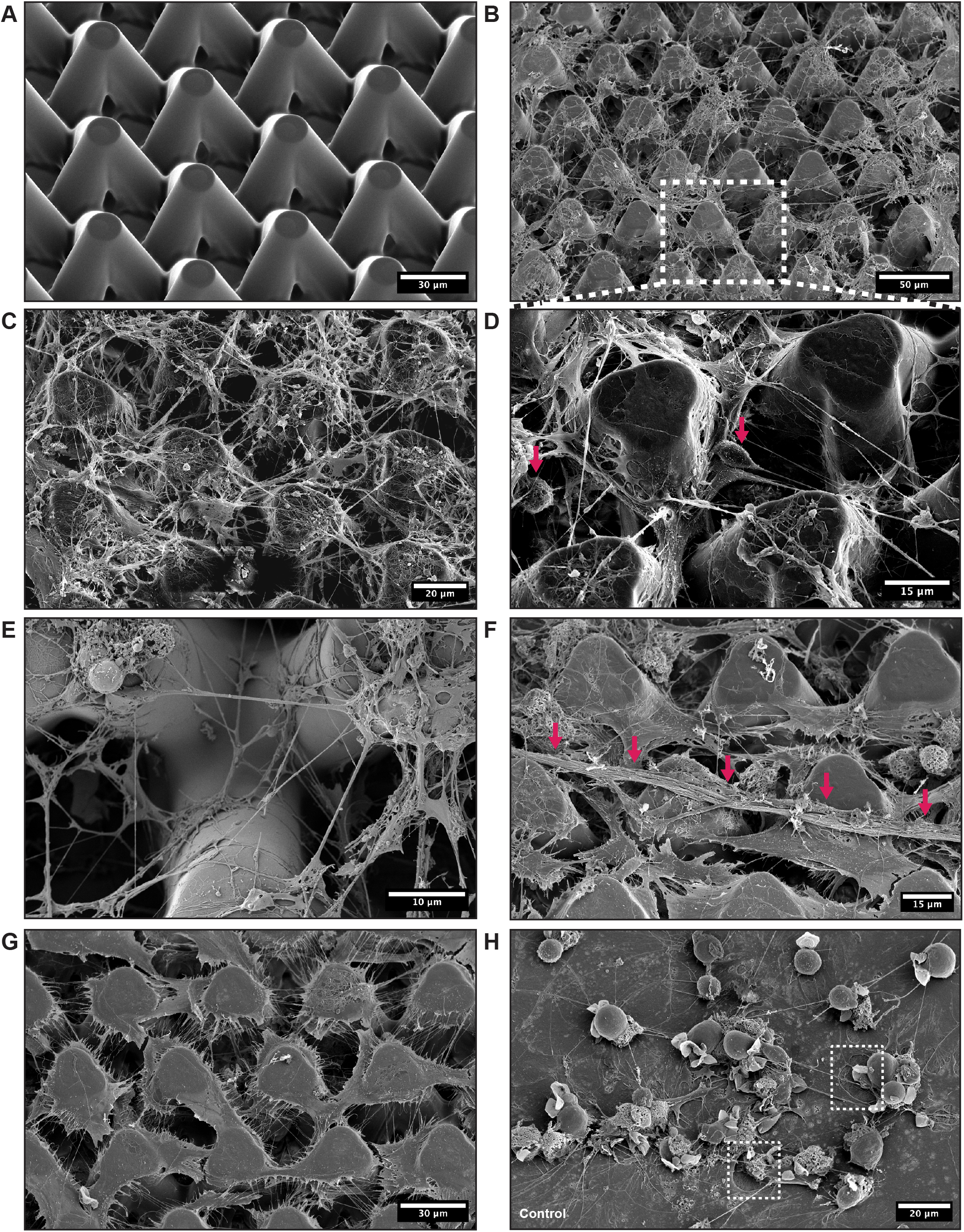
Electron microscopy of neurons growing within the 3D scaffolds. **(A.)** Tripod array before plating of neurons. **(B.)**-**(C.)** Neurons readily self-organized into 3D networks within the tripod scaffolds. **(D.)** Neuronal somata could be seen extending between the tripods, solely connected to the scaffold by their neurites. **(E.)** Neurites protruded in multiple layers, forming shortcuts between neural clusters on top of and between the tripod structures. **(F.)** Fasciculated axonal bundles were observed extending across large sections of the tripod arrays. **(G.)** Astrocytes densely populated the arrays, even forming sheets covering upper tripod regions. **(H.)** On planar controls, astrocytes formed flattened layers with neurons atop. Neurites were also observed forming dense meshes underneath the astrocytes, as indicated by the white boxes.

Astrocytes were plated two days prior to the neurons, as an astrocytic feeder layer has been shown to enhance neuronal viability (61–63) and synaptic efficacy (64). As astrocytes, unlike neurons, continue proliferation throughout the culturing period (65), extensive regions of the tripod scaffolds were covered by astrocytes by 28 DIV (**Figure 3G**). In contrast, astrocytes on planar control interfaces formed a flat monolayer across the surfaces, with neurons growing atop them (**Figure 3H**).

### Networks on Both 3D and Planar Interfaces Exhibit Structural Maturation and Plasticity at 28 DIV

Immuno-cytochemistry was employed to assess the structural maturation of the networks established on both planar interfaces and within the tripod scaffolds at 28 DIV. Growth-Associated Protein 43 (GAP43), which is highly expressed in outgrowing neurites (66, 67), was still widely expressed across the networks at this point, along with cytoskeletal markers Microtubule-Associated Protein 2 (MAP2) and *β*3-Tubulin (**Figure 4A**) (68–71). Networks on both 3D and planar interfaces also consistently exhibited high expression of the markers Neural Nuclear Protein (NeuN) and Neurofilament Heavy (NFH) (**Figure 4B**), specific to mature neurons at 28 DIV (72, 73). The presence of mature synaptic connections was furthermore confirmed by the colocalization of the pre- and postsynaptic markers synaptophysin and PSD95 (74–76). As expected, these markers were particularly abundantly expressed near neuronal somata, but were also expressed along the neurite extensions (**Figure 4C**). The high expression of both GAP43 and mature cytoskeletal and synaptic markers indicated that the networks had started to reach a mature state but were still exhibiting high levels of neural plasticity after four weeks in culture. Both control networks on planar interfaces and networks in tripod scaffolds also showed high expression of the glial marker GFAP, further indicating the high proliferation of astrocytes throughout the culturing period (77).

**Figure 4.**
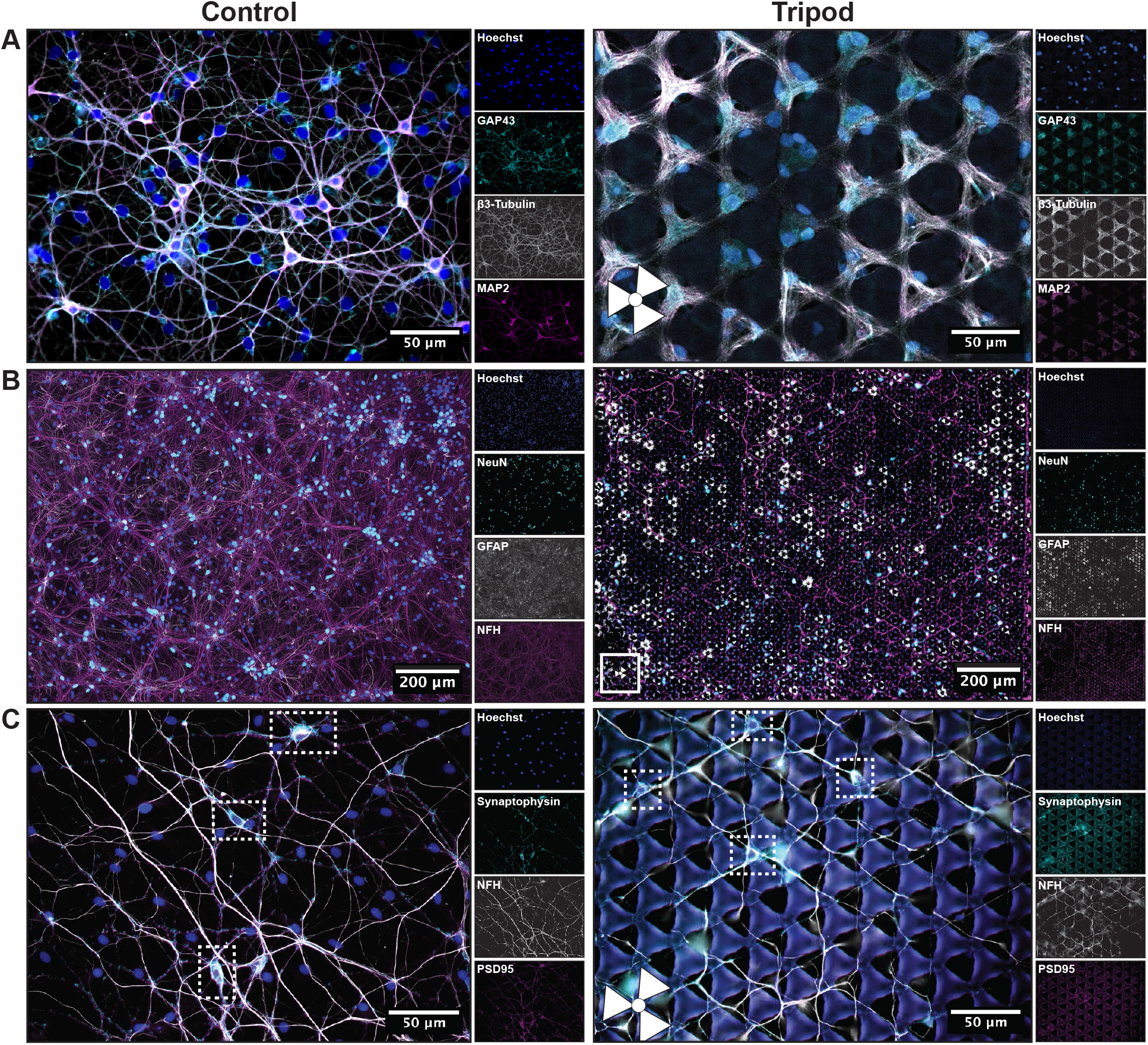
Immunocytochemistry verified structural maturity in both tripod networks and controls. **(A.)** Both network types exhibited high plasticity at 28 DIV, as indicated by the expression of GAP43, and the cytoskeletal markers Microtubule-Associated Protein 2 (MAP2) and *β*3-Tubulin. **(B.)** The high expression of Neural Nuclear Protein (NeuN) and Neurofilament Heavy (NFH), in combination with the glial marker GFAP, indicated structural maturity. **(C.)** Colocalization of the pre- and postsynaptic markers synaptophysin and PSD95 verified mature synaptic connections. A particularly high density of synapses can be seen around neuronal somata, as indicated by the white boxes. A higher exposure was required to visualize the synaptic markers compared to the other antibodies, giving rise to some autofluorescence within the tripod structures. White symbols in the lower left corner of the tripod images indicate the location of the tripods.

### Interfacing the Tripod Structures with MEAs Facilitates Functional Network Analysis

The computational capacity of the networks was investigated by interfacing the tripod structures with custom-designed nanoporous MEAs (**Figure 5A**). Gaps of 50 µm were included in the scaffold design above each electrode to ensure that the electrodes were not completely covered by tripods. Additionally, the electrodes were widened to 100 µm to increase the alignment tolerance between the microelectrodes and the gaps in the tripod design. As expected, there was some overlap between some tripods and microelectrodes. This overlap furthermore caused some deformation of the tripods in the immediate proximity to the electrodes. It also led to some platinum deposits forming in the proximity of the electrodes during electrodeposition despite SU-8’s large dielectric strength (78). However, these effects did not impact biocompatibility, nor significantly altered the available space for neurons to mature and form 3D networks.

**Figure 5.**
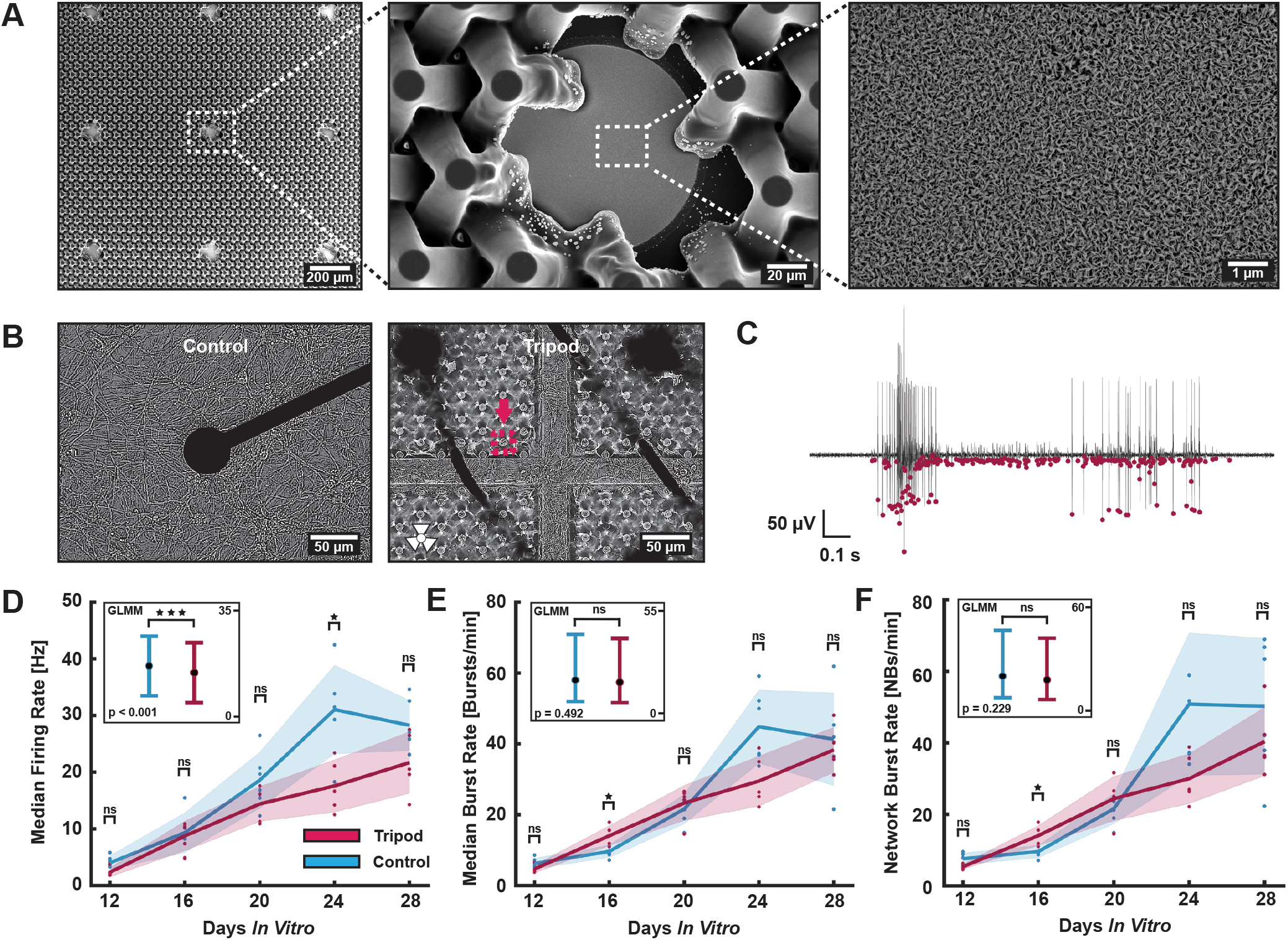
Integration of tripod structures with MEAs. **(A.)** Tripod structures were fabricated directly on top of MEAs, with nanoporous platinum electrodeposited onto the MEAs to enhance signal-to-noise ratio of the electrodes. **(B.)** Dense neural networks were established on both planar control-MEAs and tripod-MEAs. The image of the tripod network is captured at the intersection of two clamping lines, with neurites readily observed growing on top of and between the tripod structures, as outlined by the pink box. **(C.)** Networks on the tripod-MEAs exhibited a wide range of bursting and spiking dynamics, illustrated with a voltage trace showing representative bursting activity. The pink dots represent spiking events as detected by the PTSD algorithm. **(D.)** Median firing rate, **(D.)** median burst rate and **(D.)** network burst rate increased over time for both control and tripod networks. Boxes in the upper left corner of the plots show GLMM estimated group averages with 95% confidence intervals.

Neural networks were readily established on both planar MEAs and MEAs interfaced with tripod structures by 28 DIV (**Figure 5B**). From 12 DIV, both network types also exhibited complex electrophysiological activity consisting of bursting activity interspersed with individual spiking events (**Figure 5C**). Neuronal activity was detected on at least 57 out of 59 electrodes on all planar control MEAs (n = 6) and Tripod MEAs (n = 5) throughout the experimental period. Over time, the median firing rate of both the control and tripod networks gradually increased, with statistically significantly higher firing rates observed for the controls when assessing the GLMM estimated group averages (**Figure 5D**, p < 0.001). Additionally, the burst rates of individual electrodes and the network burst rates of several control networks appeared to be higher compared to the tripod networks (**Figures 5E** and **5F**, respectively). However, the large variability in the data rendered these differences statistically non-significant when assessing the GLMM estimated group averages.

### Network Integration and Entropy are Higher in Networks Established on Planar Interfaces

Information processing in the brain is characterized by a fine-tuned balance between integration and segregation of function at both the micro- and mesoscale (**Figure 6A**). This allows information to be efficiently processed in parallel in diverse neural assemblies, while also integrating these computations for more complex behaviour. Both the control and tripod networks exhibited signs of such integration-segregation dynamics in the form of strongly and more sparsely connected regions when evaluating their connectivity matrices (**Figure 6B**). Furthermore, the network coherence decreased for both control and tripod networks throughout the development, with a non-significant difference when assessing the GLMM estimated group averages (**Figure 6C**). This indicated a transition from highly synchronized bursting behaviour at 12 DIV into more complex dynamics consisting of both localized and network wide activity by the final recording at 28 DIV (**Figure 6D**). Neurons established on planar interfaces matured into more strongly connected networks compared to the controls from 20 DIV (**Figure 6E**, p < 0.001). The network complexity was on the other hand only found significantly higher for the tripod networks at 16 DIV (**Figure 6F**, p = 0.003). The entropy of individual neurons was found significantly higher for the neurons in the control networks from 20 DIV (**Figure 6G**, p < 0.001), while the overall network entropy was only significantly different at 16 DIV (**Figure 6H**, p = 0.004).

**Figure 6.**
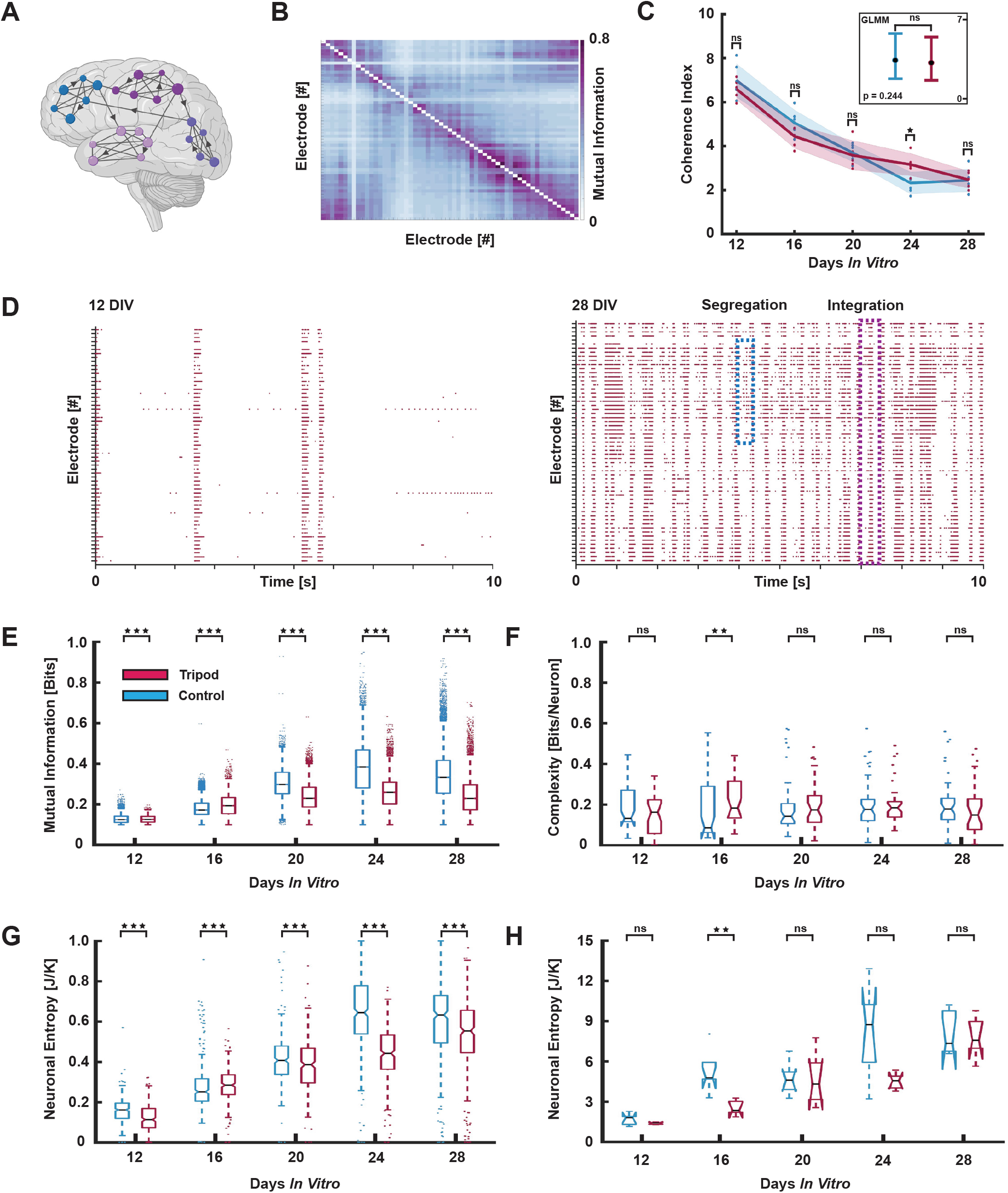
Information Theoretic Analysis of Functional Complexity. **(A.)** Information processing in the brain is characterized by both segregated and integrated functional dynamics. **(B.)** Neuronal populations on both planar control interfaces and in tripod scaffolds evolved into networks with highly interconnected and more sparsely connected regions. Purple areas indicate electrodes with a high pairwise mutual information, reflecting strong connectivity, while blue areas show weaker connectivity **(C.)** The coherence index decreased comparably for both control and tripod networks during development. The box in the upper right corner shows the GLMM estimated group average with a 95% confidence interval. **(D.)** Networks on planar control interfaces and within tripod scaffolds transitioned from sparse but highly synchronous activity during development to more complex dynamics with both synchronized and desynchronized activity at 28 DIV. **(E.)** On average, pairwise mutual information was significantly higher for control networks throughout the experimental period (p < 0.001). **(F.)** Complexity was only significantly higher for tripod networks at 16 DIV (p = 0.003) **(G.)** The entropy of individual neurons was significantly higher for control networks from 20 DIV (p < 0.001). **(H.)** Total network entropy was only significantly higher for the control networks at 16 DIV (p = 0.004).

### Connectomics Reveals More Strongly Interconnected and Clustered Networks on Planar Interfaces

To further assess the computational complexity of the networks, network connectomes were analysed through the application of graph theory (**Figures 7A** and **7B**). Each electrode was treated as an individual node, and the mutual information between pairs of electrodes as their functional connectivity. The community belonging of individual nodes can be seen changing as the networks mature. However, key network components, such as hub nodes, appeared to remain constant. Nodes indicating pacemaker activity were later chosen for targeted electrical stimulations due to their high firing rate.

**Figure 7.**
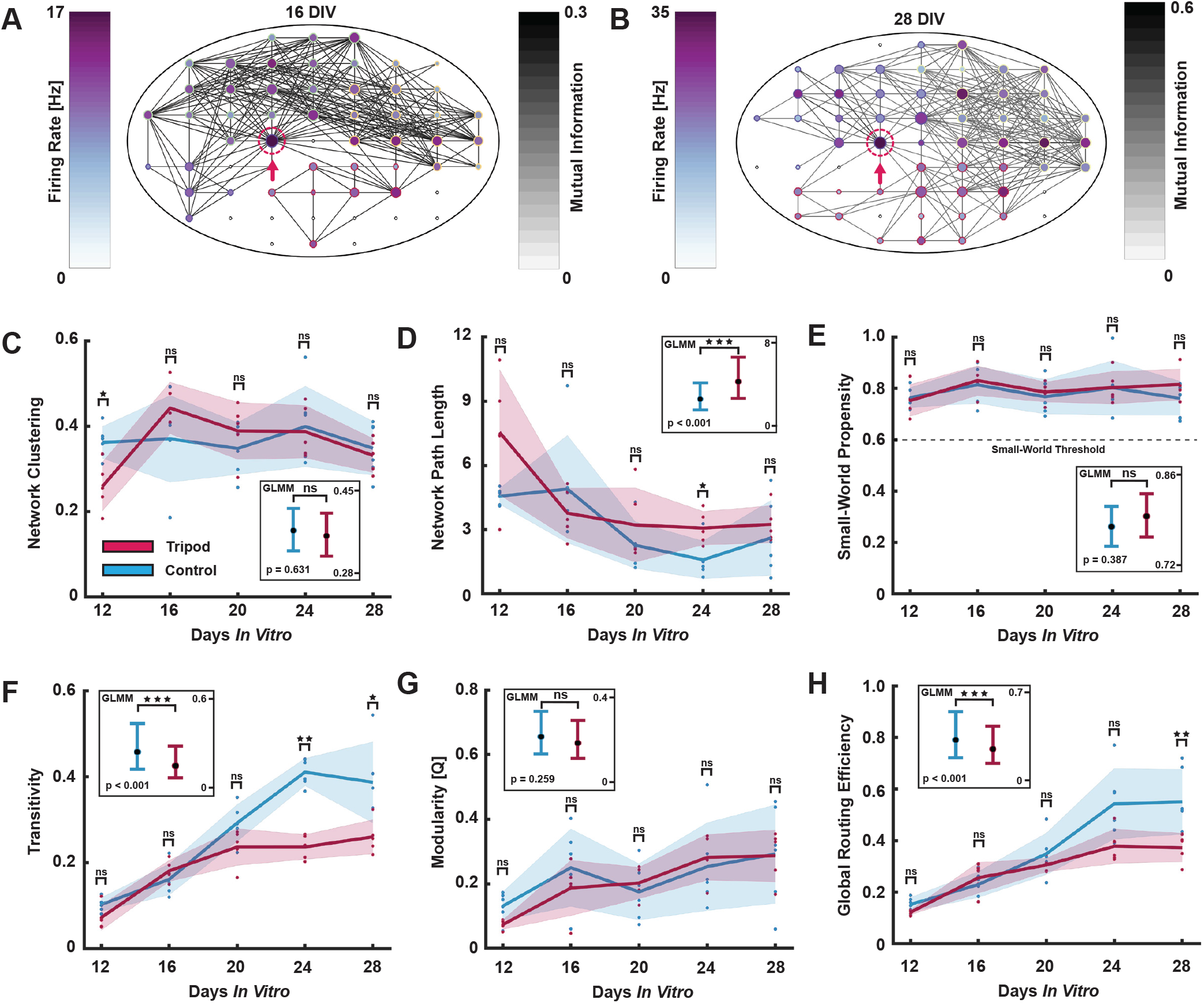
Connectomics Evaluated Using Graph Theory. **(A.)**-**(B.)** Graph representations of a tripod network at 16 DIV and 28 DIV, respectively. Node colour represents firing rate, edge colour the pairwise mutual information between nodes, node size denotes PageRank centrality, and node edge colour signifies community belonging. A pacemaker hub node is marked by the pink arrow. **(C.)** Clustering was comparable for both network types. **(D.)** The average path length was significantly higher for tripod networks. **(E.)** Both network types displayed small-world properties throughout the experimental period. **(F.)** Transitivity was significantly higher for the control networks. **(G.)** Modularity, measured using the Louvain algorithm, increased comparably over time for the two network types. **(H.)** The global routing efficiency was significantly higher for the control networks over time. The GLMM estimated group averages are shown for both network types in the corner of each plot.

At 12 DIV, the network clustering was found to be significantly higher for the controls compared to the tripod networks (**Figure 7C**, p = 0.017). However, over time, there was a non-significant difference in the GLMM estimated group averages of the two groups. The characteristic path length was on the other hand found to be significantly higher for the tripod networks compared to the controls (**Figure 7D**, p < 0.001). The small-world propensity was found comparably high for both control and tripod networks, and above the threshold of 0.6 for what is commonly considered small-world for all networks (**Figure 7E**). Here, the difference in the GLMM estimated group averages was non-significant. Transitivity, which measures the probability that adjacent nodes are interconnected, increased at a comparable rate for both network types until 20 DIV (**Figure 7F**), but was significantly higher for the control networks at 24 and 28 DIV (p = 0.004 and p = 0.017, respectively). This also yielded a significantly higher GLMM estimated group average for the control networks (p < 0.001). Interestingly, the modularity of the networks, signifying how easily the networks can be separated into distinct communities, increased comparably over time for both control and tripod networks (**Figure 7G**). The global routing efficiency, on the other hand, followed a similar trend as the transitivity and became significantly higher for the control networks at 28 DIV (p = 0.009). The GLMM estimated group average of the routing efficiency was also found significantly higher for the controls (**Figure 7H**, p < 0.001).

### Stimulations Induce Clear Responses in Networks on Both Planar Interfaces and in Tripod Scaffolds, but Impact Network Entropy Differently

After the final recording of spontaneous activity at 28 DIV, electrical stimulations were administered to assess the responsiveness and impact of stimulation on activity within the control and tripod networks. Two distinct stimulation protocols were applied, as detailed in the methods section. First, a simple stimulation protocol consisting of 30 pulses with 10 ms inter spike intervals was delivered to the most active electrode in each network. This elicited discernible responses in both the control and tripod networks (**Figures 8A-8D**).

**Figure 8.**
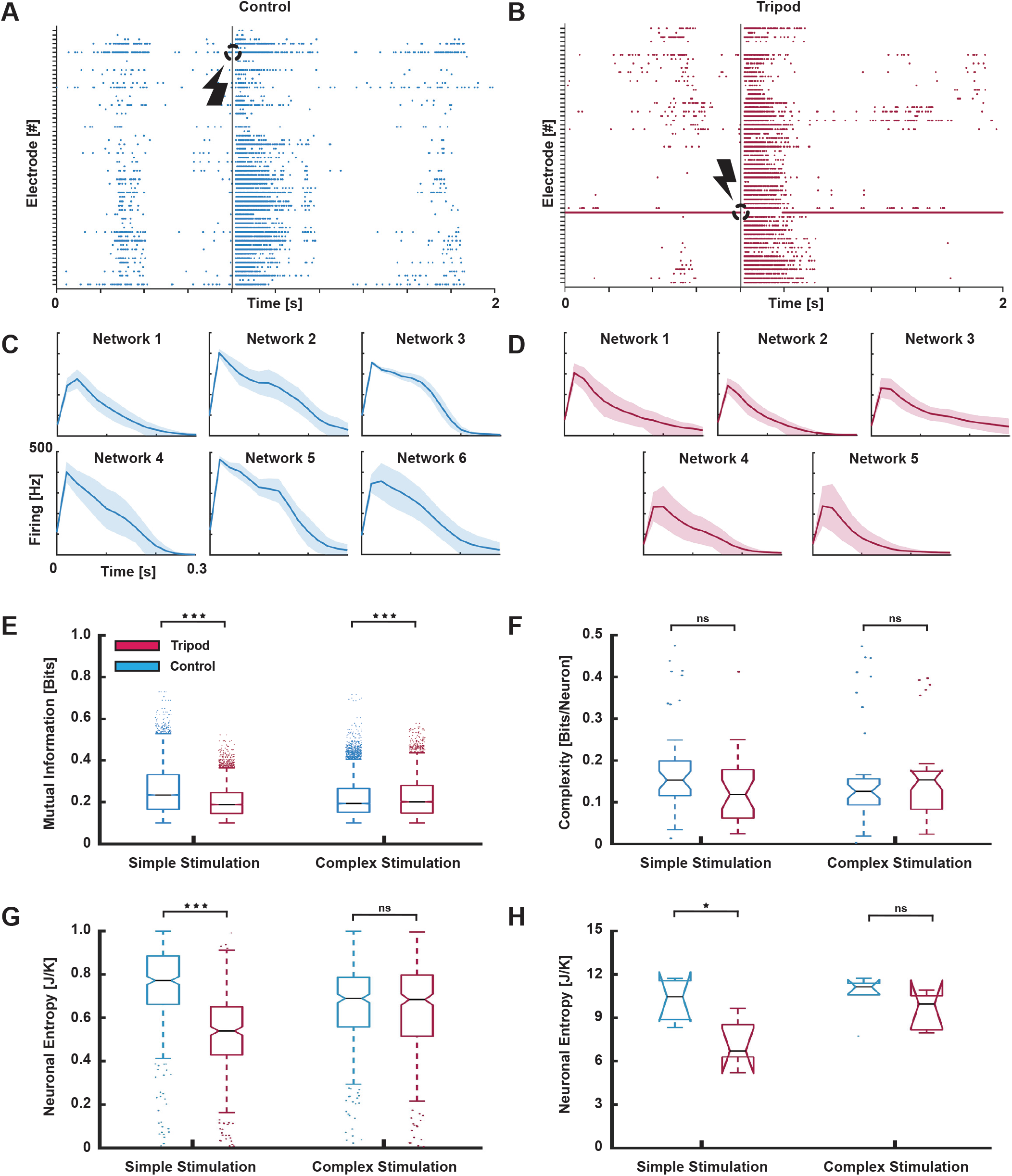
Stimulation Induced Network Responses. **(A.)**-**(B.)** Representative raster plots displaying clear responses to electrical stimulations in the control and tripod networks, respectively. **(C.)**-**(D.)** Peristimulus time histograms depicting the mean and standard deviation of responses in each control and tripod network to 30 electrical pulses with 10 ms interspike intervals. **(E.)** Functional connectivity was notably higher for the control networks in response to the simple stimulation protocol, while it was higher for the tripod networks in response to the complex stimulation protocol. **(F.)** The complexity in the electrophysiological data was comparably high for the two network types in response to both the stimulation protocols. **(G.)**-**(H.)** Individual neuron entropy and network entropy were significantly higher for the control networks in response to the simple stimulation protocol. However, the difference was non-significant for the complex stimulations, where the entropy of the tripod networks increased significantly while the entropy of the control networks remained unchanged.

Similar to the analysis of the spontaneously evoked activity, the functional connectivity strengths between pairs of electrodes were notably higher in the control networks compared to the tripod networks in response to the simple stimulation protocol (**Figure 8E**, p < 0.001). Conversely, in response to the complex stimulation protocol, connectivity was significantly higher in the tripod networks (p < 0.001). Nevertheless, complexity remained comparably high for the two network types (**Figure 8F**). Neuronal entropy was also greater in the control networks than in the tripod networks for the simple stimulation protocol (**Figures 8G** and **8H**, p > 0.001 and p = 0.0173, respectively). However, for the complex stimulation protocol, both the entropy of individual neurons and the networks established in tripod scaffolds increased to levels comparable to those of the controls.

## Discussion

Several challenges had to be addressed during optimization of the different fabrication steps. One significant issue was the recurrent delamination of the tripods from the substrate, attributed to the large differences in the thermal expansion coefficients of SU-8 and borosilicate/Si_3_N_4_, as well as the limited contact area between the tripod legs and the substrate. While slower PEBs and the introduction of clamping lines partly mitigated this issue, utilizing a collimated UV lamp could offer a more uniform exposure of the photoresist in future studies. Additionally, achieving uniform deposits of nanostructured platinum across the microelectrode posed challenges. The presence of the tripod legs during electrodepositions led to unstable growth of platinum islands on the electrodes, likely due to alterations in diffusion kinetics of chloroplatinic acid towards the interface. This issue was particularly pronounced for depositions above 3 min. Nevertheless, even a brief deposition of porous platinum has been shown to significantly improve the signal-to-noise ratio of microelectrodes (5, 79), and was therefore deemed suitable for improving the electrode properties significantly without adverse effects.

The presented fabrication approach offers notable advantages over previous studies, demonstrating high reproducibility, optical transparency, and compatibility with electrophysiological recording techniques. Additionally, the straightforward integration of PDMS chambers as cell culturing compartments onto the substrates surrounding the SU-8 structures for the tripod-MEAs indicates compatibility of these 3D structures with PDMS-based microfluidics (80, 81). One advantage with keeping the SU-8 structures below 100 µm is also the ensured accessibility to the entire neural network both visually and electrophysiologically throughout experiments, given the microelectrodes’ capability to detect signals up to ∼ 100 µm away from neurons (82). Future improvements could however include implementation of 3D electrodes on these interfaces for enhanced signal acquisition from the upper layers of cells (19). Alternative strategies could involve the use of 3D electrodes on external probes lowered into the cultures (83), or introducing dopants into the SU-8 to confer electrical conductivity to the tripods themselves (84, 85).

Combining advanced engineering approaches such as 3D scaffolds and microfluidics can contribute further towards more biologically relevant engineered model systems, enhancing the neural networks’ ability to mature into computational entities exhibiting attributes of dynamical richness and computational complexity (86–88). Modular and topologically complex structures can potentially enhance the functional capacity and input representation of networks (89), and contribute to recapitulation of the complex dynamics of cortical information processing in the brain (90–92). In line with this, the neuronal cell populations plated on both planar controls and in tripod scaffolds evolved into networks with hallmarks of high computational capacity, such as a modular architecture, small-world properties and a balance between both segregated and integrated functional dynamics. A key finding was however that the functional connectivity, measured as pairwise mutual information, was consistently higher for the networks established on planar surfaces throughout the experimental period. Additionally, control networks displayed higher firing and burst rates, more efficient global signal transfer, and greater network clustering. These observations align with previous studies, suggesting lower synchronization levels in 3D networks compared to those on planar surfaces (32).

When considering network synchronization and integration, it is important to acknowledge that engineered networks commonly lack myelination, a key factor in synchronizing distant neuronal populations in the brain. Research indicates that demyelination of axons can lead to energy depletion, resulting in increased periaxonal potassium levels and ephaptic coupling, leading to hyperexcitability (93–95). Given that networks formed on planar surfaces develop dense, overlapping, and unmyelinated neurite meshworks, ephaptic coupling may contribute to enhanced activity. This effect may be further exacerbated by the dense monolayer of astrocytes covering up the neurite mesh, as observed in this study. Conversely, in 3D networks, neurites can extend and interconnect across multiple layers, reducing the overlap between neurite protrusions and potentially mitigate the impact of ephaptic coupling on network excitability. Moreover, astrocytes in the tripod networks were observed to envelope large portions of the arrays, likely providing neurons with nutritional and ionic buffering support from various directions. This phenomenon could have significant implications for network homeostasis and activity, representing an intriguing avenue for future research in leveraging these scaffolds.

Another intriguing observation was the higher entropy exhibited by the control networks compared to the tripod networks, both during spontaneously evoked activity and in response to the simple stimulation protocol. Neuronal entropy is a crucial network feature, characterizing rich dynamics and neuronal adaptation in the brain (96). Furthermore, it has previously been found to negatively correlate with functional connectivity strength, suggesting that weaker connections may enable more diverse repertoires of network dynamics (97). On the other hand, excessively high entropy may diminish the information gain from the system (29). In the case of networks on planar interfaces, the strong integration may limit the networks’ ability for information gating, thus increasing their entropy. Conversely, the 3D networks can establish connections in the z-direction, potentially creating shortcuts that contribute to more complex information gating between neural clusters in different scaffold regions. While the complexity of both control and tripod networks appeared comparably high, the underlying features contributing to this complexity differed, suggesting that topological dimensionality influences the balance between integrated and functionally segregated dynamics in networks. Interestingly, the tripod networks exhibited significantly stronger functional integration in response to the complex stimulation protocol, with entropy rising to a level comparable to that of control networks. This variation in stimulation susceptibility underscores the intricate interplay between dimensionality, plasticity, spontaneously evoked activity, and external input in shaping neural network dynamics.

## Conclusion

The topology of neural networks profoundly influences information processing, transfer, and storage in the brain. However, most *in vitro* studies are conducted with neural networks established on planar interfaces, lacking a 3D microenvironment for the networks to self-organize in. In this study, we introduced a novel method for fabricating 3D SU-8 scaffolds compatible with neuronal growth using inclined photolithography. Theses interfaces are compatible with a range of techniques for studying the networks’ structure-function dynamics, including MEAs. Through information- and graph theoretic analyses, we showcased that neurons within these scaffolds develop into networks with high computational capacity, exhibiting complex, spontaneously evoked activity and consistent responses to electrical stimulation. Comparison with networks on planar interfaces revealed a similarly complex interplay between segregated and integrated information processing. Key differences were however that the control networks displayed significantly stronger functional connectivity, measured using the electrodes’ pairwise mutual information, and higher overall entropy. In summary, by better recapitulating the 3D topology of neural networks in the brain, these interfaces can be used to advance our understanding of how the topology of neural networks impact network information processing and computations in both health and disease.

## AUTHOR CONTRIBUTIONS

The author contributions follow the CRediT system. **NWH**: Conceptualization, Methodology, Software, Investigation (chip design & manufacturing, cell experiments, ICC, electrophysiology, bioSEM imaging, formal analysis), Writing – Original Draft, Visualization. **KGK**: Methodology, Investigation (chip design & manufacturing, confocal microscopy, SEM imaging), Writing – Review & Editing. **ASN**: Methodology, Investigation (stage design, chip design & manufacturing, SEM imaging, contact angle measurements), Writing – Review & Editing. **PS, AS, IS**: Conceptualization, Methodology, Writing – Review & Editing, Resources, Funding Acquisition.

## FUNDING

NTNU Enabling technologies and the Central Norway Regional Health Authority are acknowledged for funding this research. The Research Council of Norway is acknowledged for the support to the Norwegian Micro- and Nano-Fabrication Facility, NorFab, project number 295864.

## ACKNOWLEDGEMENTS

We would like to thank senior engineer Astrid Bjørkøy and the Center for Advanced Microscopy (CAM) at the Department of Physics, Faculty of Natural Sciences, NTNU for technical assistance and access to confocal microscopy infrastructure. We also thank engineers Mark Chiappa, Martijn de Roosz, Jakob Vinje and Mathilde Barriet at NTNU NanoLab for technical assistance. Furthermore, we acknowledge Nan Tostrup Skogaker for training and access to the electron microscopy core facility, NTNU. Åste Brune Tomren and Polina Malahov are acknowledged for help with experimental cell work during pilot studies when developing the tripod technology. We would also like to thank Prof. Michela Chiappalone and Prof. Sergio Martinoia, University of Genova for generously providing the scripts for the Precise Timing Spike Detection algorithm and the logISI burst detection.

## COMPETING FINANCIAL INTERESTS

The authors declare that the research was conducted in the absence of any commercial or financial relationships that could be construed as a potential conflict of interest.

## Notes

### Competing Interest Statement

The authors have declared no competing interest.

### Summary of Updates

Figure 1 has been updated to improve image quality.

