## Supplementary Materials for "Functional Complexity of Engineered Neural Networks Self-Organized on Novel 3D Interfaces"

**Nicolai Winter-Hjelm[1,✉], Kasper Grøndahl Klausen[2], Amund Stensrud Normann[2], Axel Sandvig[1,3,4,5], Ioanna Sandvig*[1,✉], and Pawel Sikorski*[2,✉]**

[1]Department of Neuromedicine and Movement Science, Faculty of Medicine and Health Sciences, Norwegian University of Science and Technology (NTNU), Norway
[2]Department of Physics, Faculty of Natural Sciences, Norwegian University of Science and Technology (NTNU), Trondheim, Norway
[3]Department of Neurology and Clinical Neurophysiology, St Olav´s University Hospital, Trondheim, Norway
[4]Department of Community Medicine and Rehabilitation, Umeå University Hospital, Umeå, Sweden
[5]Department Neuro, Head and Neck, Section for Spinal cord and Head Injuries, Umeå University Hospital, Umeå, Sweden

****These authors have contributed equally to this work.***

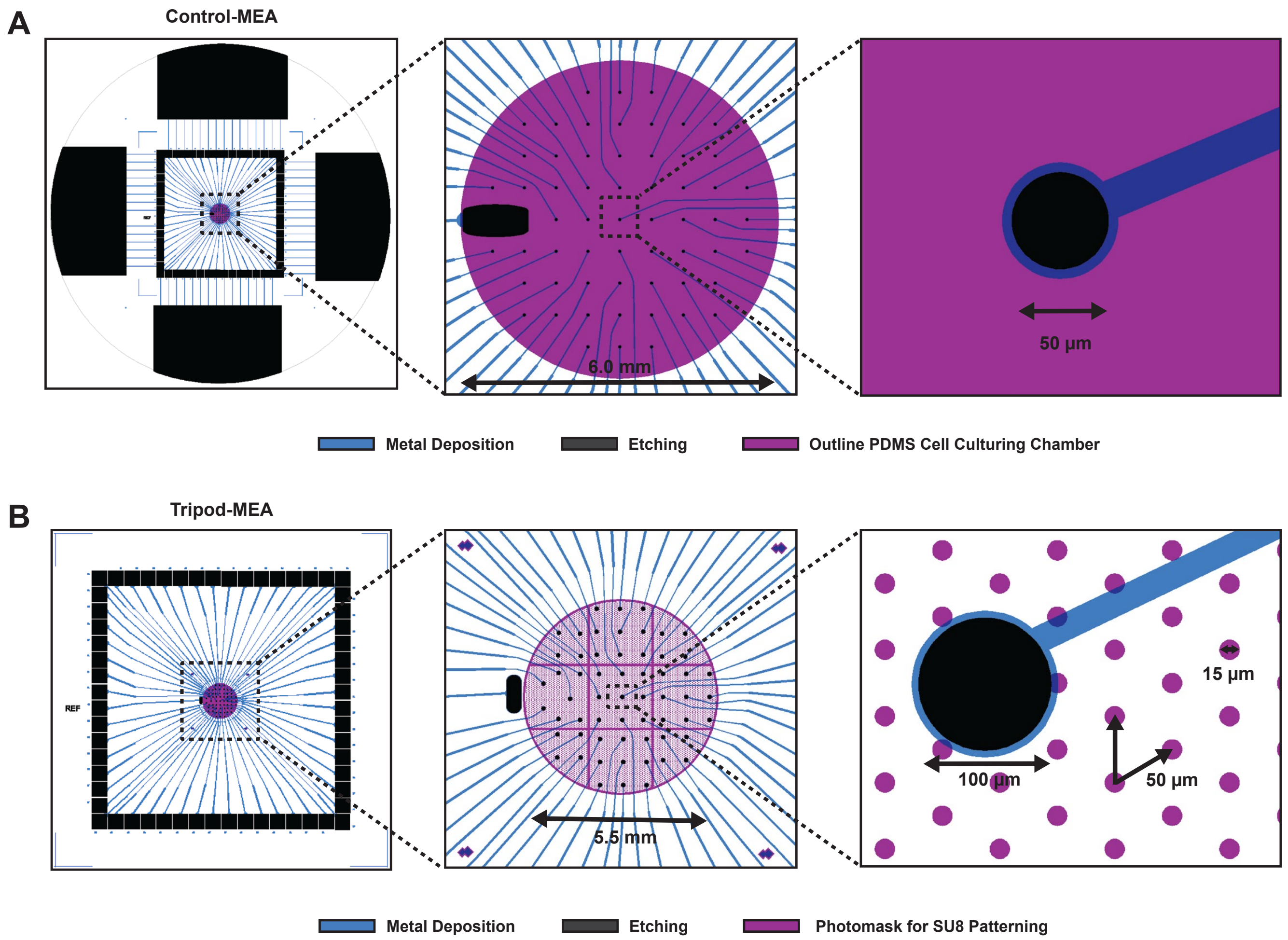


**Figure S1 | CAD designs for the control- and tripod-MEAs. (A.)** Design used to fabricate the control-MEAs. The four large contact pads along the edges of the wafer were used for connecting the MEAs to the potentiostat during depositions of nanoporous platinum. Following depositions, the wafers were subsequently cut into 49 x 49 mm squares with a wafer saw along the four corner marks, hence disconnecting the 60 smaller contact pads from the larger ones. The third layer, *outline for the PDMS cell culturing chamber*, is only included to illustrate the outline of the PDMS chambers. **(B.)** Design used to fabricate the tripod-MEAs. The *photomask for SU-8 patterning* consisted of $15\,\mu m$ apertures in a hexagonal grid pattern with $50\,\mu m$ spacing. Empty spaces of $50\,\mu m$ diameter were included at the positions of the electrodes to avoid tripods covering up the electrodes. The electrodes were $100\,\mu m$ in diameter to increase alignment tolerance. However, it is worth noting that only $50\,\mu m$ of the electrodes would be fully available to the neurons, similar as for the controls, as tripods would cover up parts of them. $50\,\mu m$ diameter clamping lines were included with an interspacing of $1.5\,mm$, as well as around the outer edge of the design to prevent heat folding and delamination of the SU-8 from the substrate. The SU-8 pattern was made $5.5\,mm$ in diameter to avoid overlap between the SU-8 and PDMS during bonding of the cell culturing chambers.

### Photomask Fabrication

Mask designs for the tripod structures were made using Clewin 4. A custom-made Python script was furthermore used to generate large hexagonal dot patterns. A script to create dot patterns of customizable size and distance can be found at https://github.com/amundsno/CleWin-Hexagonal-Dot-Array-Generator.

$1\,\mathrm{mm}$ thick 4-inch borosilicate wafers ($100\,\mathrm{mm}$ Borofloat33, Plan Optik) were used as substrates for the photomasks. The wafers were washed subsequently in acetone and IPA for $1\,\mathrm{min}$ each to remove organic contaminants, before being plasma cleaned for $2\,\mathrm{min}$ in $200\,\mathrm{sccm}$ $O_2$ plasma at $40\,\mathrm{kHz}$ generator frequency. A $3\,\mathrm{min}$ dehydration bake was conducted at $150\,°\mathrm{C}$. Following cleaning, $100\,\mathrm{nm}$ chromium was deposited onto the substrate at $5\,\mathrm{Å\,s^{-1}}$ while rotating the sample for uniform deposition (E-beam Evaporator, Pfeiffer Vacuum Classic 500). The photoresist SPR700-1.0 (Dow Europe GmbH) was spin coated onto the substrates at $3000\,\mathrm{rpm}$ for $30\,\mathrm{s}$ at $1000\,\mathrm{rpm/min}$ and soft-baked at $95\,°\mathrm{C}$ for $1\,\mathrm{min}$ to form a protective $1\,\mathrm{\mu m}$ layer. Subsequently, the wafer was diced into 12x12 mm squares using a wafer saw (DAD323, DISCO).

Following dicing, the wafers were cleaned, and a new layer of $1\,\mathrm{\mu m}$ thick SPR700 was spun onto the substrate. A maskless aligner (MLA150, Heidelberg) was used to transfer the hexagonal dot pattern onto the resist with a $405\,\mathrm{nm}$ laser at $50\,\mathrm{mJ/cm^2}$. A post exposure bake was conducted at $115\,°\mathrm{C}$ for $1\,\mathrm{min}$, before developing the resist in ma-D332/s (micro resist Technology GmbH) for $50\,\mathrm{s}$ and thoroughly rinsing in DI water. Next, the substrates were descummed in $100\,\mathrm{sccm}$ $O_2$ plasma for $1\,\mathrm{min}$ at $20\,\mathrm{kHz}$ generator frequency. The substrate was subsequently etched using Chromium Etchant Standard solution (Sigma-Aldrich, 651826).

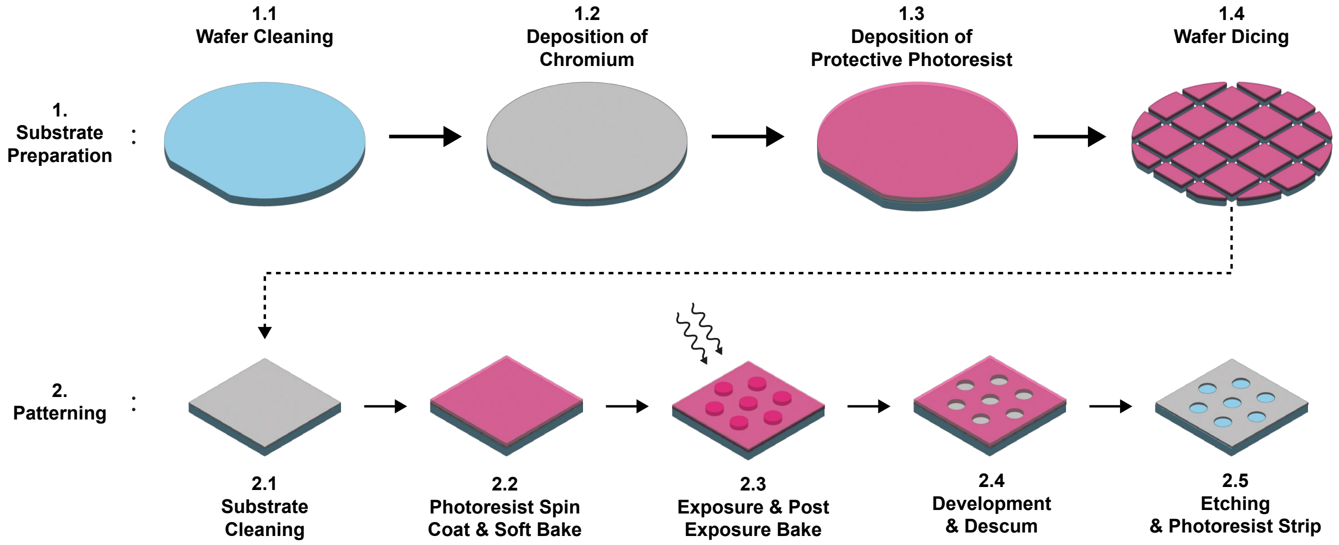


**Figure S2 | Illustration of the photomask fabrication steps.** A polished $1\,\mathrm{mm}$ thick 4" glass wafer was cleaned **(1.1)**, and a $100\,\mathrm{nm}$ chromium layer deposited onto the substrate **(1.2)**. A protective photoresist layer was subsequently spin coated onto the substrate **(1.3)**, and the substrate diced into 12x12 mm samples **(1.4)**. Next, the substrate was cleaned **(2.1)**, and a new layer of photoresist was spin coated onto the substrate **(2.2)**. The photoresist was exposed using a maskless aligner according to the hexagonal dot pattern and post exposure baked **(2.3)**. Next, developing was performed, and the substrate descummed using oxygen plasma **(2.4)**. Eventually, chemical etching was conducted to imprint the design onto the chromium layer and finalize the photomask **(2.5)**.

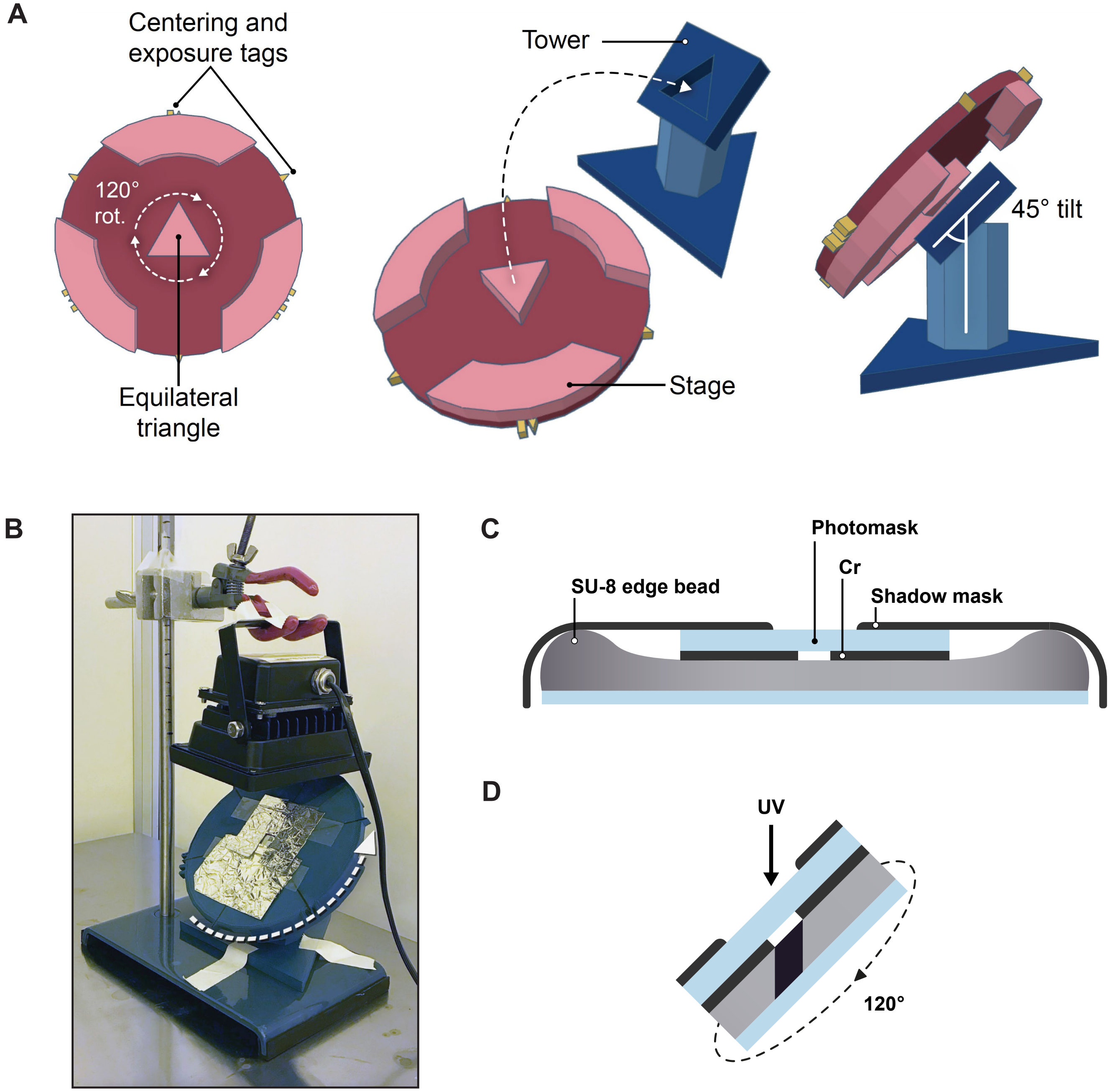


**Figure S3** | **Setup for inclined lithography exposures. (A.)** Design of the substrate holder used during exposures, made in TinkerCAD. The holder consisted of two parts: A stage and a tower. The stage had a $45^\circ$ tilt when mounted to the tower. Furthermore, the stage was mounted to the tower with an equilateral triangle, to facilitate $120^\circ$ rotations between exposures. Additionally, centering and exposure tags were included to enable accurate centering of the sample and to keep track of the number of exposures performed. **(B.)** After mounting the sample to the stage, and thereafter the stage to the tower, exposures were conducted with a $10\,\mathrm{W}$ XT004 UV curing lamp from Extral Light Store. The UV lamp was mounted above the sample at $5\,\mathrm{cm}$ distance using a clamp stand. **(C.)** The photomask was pushed into contact with the sample with the chromium layer facing down. Due to the edge bead of the SU-8, it was essential that the mask was slightly smaller than the substrate so that the mask could come into proper contact with the photoresist. Furthermore, aluminium foil was used to cover up the edges of the photoresist not protected by the photomask to avoid unintentional crosslinking of the SU-8. **(D.)** Each exposure lasted $25\,\mathrm{s}$, yielding a total dosage of $525\,\mathrm{mW/cm^2}$. Each sample was furthermore exposed three times, with a $120^\circ$ rotation in between each exposure to create the tripod structures seen in the main paper.
